# Small non-coding RNA Interactome Capture reveals pervasive, carbon source-dependent tRNA engagement of yeast glycolytic enzymes

**DOI:** 10.1101/2022.07.14.500110

**Authors:** Claudio Asencio, Thomas Schwarzl, Sudeep Sahadevan, Matthias W. Hentze

## Abstract

Small non-coding RNAs fulfill key functions in cellular and organismal biology, typically working in concert with RNA-binding proteins (RBPs). While proteome-wide methodologies have enormously expanded the repertoire of known RBPs, these methods do not distinguish RBPs binding to small non-coding RNAs from the rest. To specifically identify this relevant subclass of RBPs, we developed small non-coding RNA interactome capture (snRIC_2C_) based on the differential RNA-binding capacity of silica matrices (2C). We define the *S. cerevisiae* proteome of nearly 300 proteins that specifically binds to RNAs smaller than 200 nucleotides in length (snRBPs), identifying informative distinctions from the total RNA-binding proteome determined in parallel. Strikingly, the snRBPs include most glycolytic enzymes from yeast. With further methodological developments using silica matrices, 12 tRNAs were identified as specific binders of the glycolytic enzyme GAPDH. We show that tRNA engagement of GAPDH is carbon source-dependent and regulated by the RNA polymerase III repressor Maf1, suggesting a regulatory interaction between glycolysis and RNA polymerase III activity. We conclude that snRIC_2C_ and other 2C-derived methods greatly facilitate the study of RBPs, revealing previously unrecognised interactions.

## Introduction

RNA-protein interactions govern not only gene expression from beginning to end, but are also crucial for the function of cellular machines such as the ribosome, the spliceosome, the signal recognition particle, telomerase and many others (Glisovic et al.2008, Mitchell & Parker2014, Gerstberger et al.2014, Singh et al.2015). Recent work suggests that the biological scope of RNA-protein interactions and RNA-binding proteins (RBPs) is even larger than previously anticipated (Baltz et al.2012, Castello et al.2012). Against this background, the development of methods to study RNA-protein interactions and RBPs promises fundamental new insights.

Decades ago, UV photo-crosslinking was shown to catalyse covalent bond formation between RNA and proteins at zero distance (Brimacombe et al.1988, Hockensmith et al.1986) without similarly promoting protein-protein crosslinking (Greenberg1979, Pashev et al.1991, Suchanek et al.2005). To identify the poly(A) RNA-binding proteome of cultured cells comprehensively, RNA interactome capture (RIC) combined UV photo-crosslinking with oligo-dT capture of polyadenylated RNAs followed by mass spectrometry (Baltz et al.2012, Castello et al.2012). RIC was subsequently refined by introduction of locked nucleic acid (LNA)-modified dT-capture probes (enhanced RIC, eRIC), allowing more stringent conditions and improving the signal to noise ratio of conventional RIC (Perez-Perri et al.2021, Perez-Perri et al.2018). However, both RIC and eRIC only identify RBPs that bind polyadenylated RNAs. To identify RBPs irrespective of the class of RNAs that they bind to, OOPS (Queiroz et al.2019) and XRNAX (Trendel et al.2019) were developed to extract crosslinked RBPs from the interphase between aqueous and organic solvents after a phenol extraction of RNA, while the PTex method uses two organic solvents (Urdaneta et al.2019).

We (Asencio et al.2018) and others (Shchepachev et al.2019) recently reported that the silica matrices commonly used for total RNA isolation also retain RBPs that are covalently crosslinked to RNA, a method we refer to as complex capture (2C). Here, we explored 2C for the determination of the total RNA-binding proteome of the yeast *Saccharomyces cerevisiae*, establishing RIC_2C_. Unexpected observations then motivated the development of further downstream applications of 2C, especially for the identification of the cellular RNAs that bind to an RBP of interest (CLIP_2C_) and the determination of those RBPs that bind to small non-coding RNAs (snRIC_2C_), a polyfunctional class of RNAs with multiple regulatory roles. These technical advances led us to uncover a novel connection between tRNAs, glycolytic enzymes and carbon metabolism in yeast.

## Results

### RIC_2C_ identifies 983 RBPs in yeast

We recently reported that commercially available silica columns used to purify RNA can also be employed for the co-purification of crosslinked RBPs to capture covalently linked RNA-protein complexes, called complex capture (2C)(Asencio et al.2018). To utilize 2C for the determination of a high confidence RNA-binding proteome of the yeast *Saccharomyces cerevisiae*, we irradiated cultured cells with 3 J/cm^2^ UV light at 254 nm, using non-irradiated cells as negative controls. Lysates from three independent biological replicates were subjected to a first round of 2C purification. To minimise residual contamination with DNA-binding proteins, we treated the 2C eluates with DNase I and conducted a second round of 2C. Subsequently, RBPs in the second round eluates were released by RNase I treatment, TMT-labelled, and analyzed by mass spectrometry (Figure 1A). The enrichment of RBPs by RIC_2C_ compared to the negative controls is strong and highly consistent (Suppl. Figure 1), identifying 983 RBPs from yeast (Figure 1B, table 1). Gene Ontology (GO) term enrichment analysis confirms ribosomal proteins as a highly enriched category in the crosslinked samples, reflecting that RIC_2C_ is not restricted to mRNA-binding proteins and efficiently captures the total RNA binding proteome (Figure 1C).

**Figure 1.**
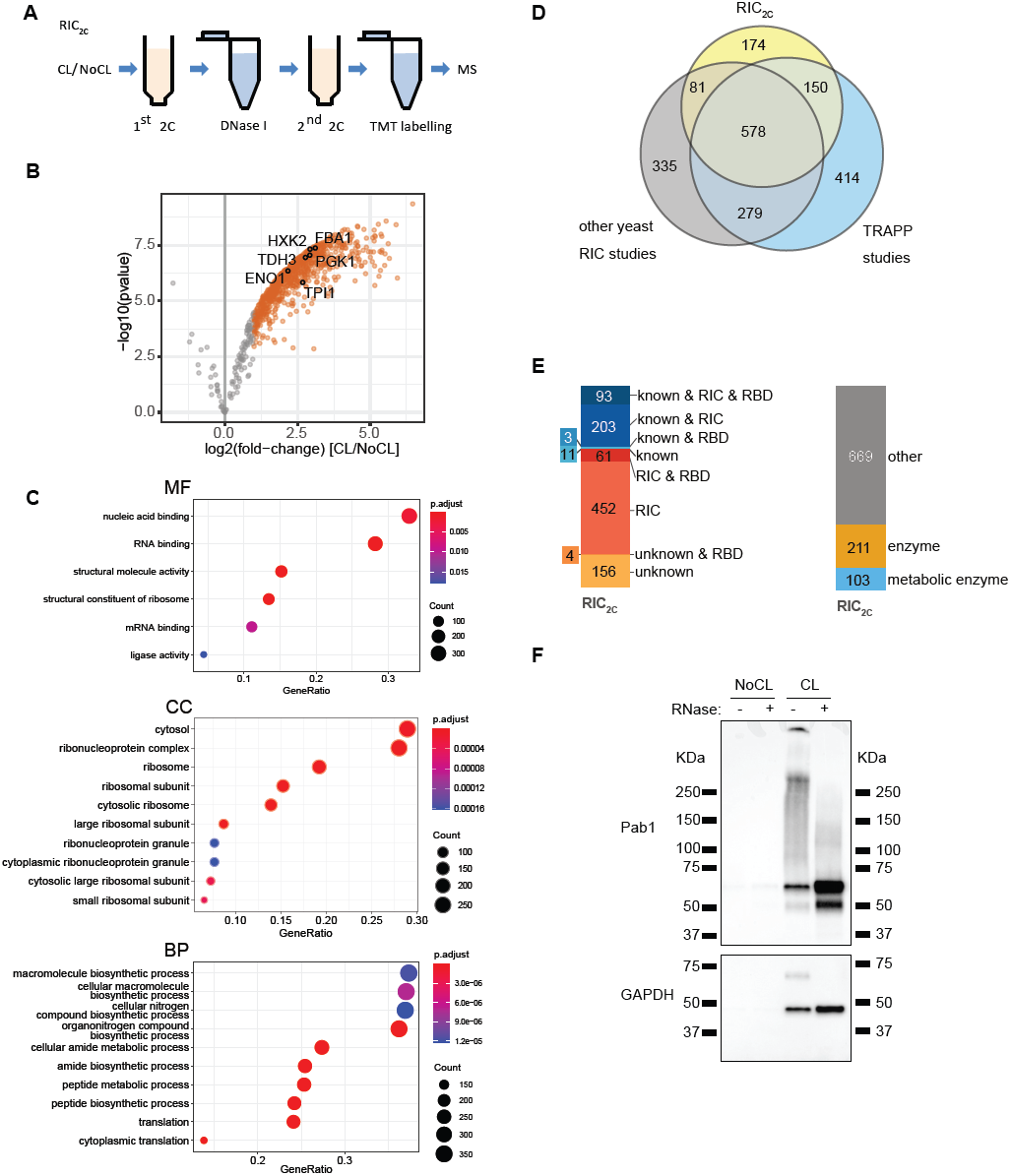
2C total RNA interactome capture (RIC_2C_) identified 983 RBPs in yeast. **A)** Schematic representation of RIC_2C_. UV-crosslinked and non-irradiated negative controls were subjected to a first round of 2C. Any residual DNA in the eluates was digested by DNase I and the RNA and RNA-protein adducts were re-purified by a second 2C extraction. Eluates from the second round were RNase I treated and proteins were subjected to TMT labelling and mass spectrometry analysis. **B)** Volcano plot displaying Log2 fold change of protein abundance vs –Log10 p-value after RIC_2C_ of CL and NoCL samples. Grey dots represent proteins displaying no statistically significant difference. Orange dots represent proteins statistically enriched (logFC≥1 and p-value≤0.05) in CL over NoCL samples. Glycolytic enzymes statistically enriched in the CL fraction are highlighted with black circles. **C)** GO-term analysis of RBPs identified by RIC_2C_. **D)** Venn diagram showing the overlap between the RBPs detected by RIC_2C_, TRAPP (Shchepachev et al.2019) and a compendium of other RNA interactome capture experiments in yeast (Beckmann et al.2015, Brannan et al.2016, Kramer et al.2014, Matia-Gonzalez et al.2015, Mitchell et al.2013, Ray et al.2013, Scherrer et al.2010, Shchepachev et al.2019, Tsvetanova et al.2010). **E)** Analysis of the RBPs detected after RIC_2C_ in yeast. RBPs were categorized according to experimental evidence described in literature (“known”), their detection on RIC experiments (RIC) or content of RNA binding domains (RBD). Novel RBPs detected by RIC_2C_ unrelated to previous experimental evidence and not detected on any RIC experiment were categorized as “unknown”. **F)** Validation of two RBPs by 2C-WB. 10ug of 2C RNA from CL and NoCL samples were treated or not with RNase I, separated by SDS-PAGE, blotted to a nitrocellulose membrane and probed against Pab1 and GAPDH antibodies.

Comparison with published datasets of RBPs in yeast suggests that the repertoire of RBPs in yeast may be nearing completion, with 174 additional novel RBPs (Figure 1D) identified in our dataset, including 156 RBPs without known RNA-binding domain (RBD) (Figure 1E). Like previous studies, RIC_2C_ identified numerous metabolic enzymes, including several members of the glycolytic pathway, as RBPs (Figure 1B, 1E & table 1).

To validate RIC_2C_, eluates from crosslinked and non-crosslinked samples were treated with RNase I or left untreated, and subjected to immunoblotting for the RBPs Pab1 and GAPDH (Tdh3), respectively. Both proteins were only retained by RIC_2C_ after crosslinking. Furthermore, Pab1 showed the characteristic RNA-induced smear that collapses after RNase treatment into more defined bands just exceeding the molecular mass of the native protein (Figure 1F, top panel). Interestingly, crosslinked GAPDH shows a far more defined band of slower migration rather than a smear without RNase treatment, which again collapses into a faster migrating GAPDH band after RNA digestion (Figure 1F, bottom panel). GAPDH has been found to bind tRNAs in HeLa cells (Singh & Green1993), and our results reveal that yeast GAPDH apparently binds a relatively homogenous class of low molecular mass RNAs. These results suggest that yeast GAPDH may also bind tRNAs and exemplify the utility of RIC_2C_.

### CLIP_2C_ identifies several tRNAs as specific GAPDH-binding partners

To follow up on the unexpected GAPDH result, we decided to determine its RNA binding partners. Existing CLIP protocols have been very successful with canonical RBPs like hnRNPs, splicing factors or Pab1 (Baejen et al.2014), but show limitations with non-canonical RBPs, where often only a minor fraction of the cellular protein is bound to RNA. Such a situation typically causes signal to noise issues from high background.

We reasoned that enrichment of the RNA-bound fraction by 2C before immunoprecipitation could help to address this situation. To test this notion, we compared GAPDH immunoprecipitations from input lysates and 2C eluates and evaluated the efficiencies of capturing crosslinked GAPDH-RNA complexes. Input and unbound fractions, together with eluates from the IPs were analyzed by western blotting. While the IP from the input lysate showed a stronger overall signal, including more background above and below the expected size, the pattern was unaffected by RNase treatment, strongly suggesting that most of the immunoprecipitated protein was not bound to RNA (Suppl. Figure 2). By contrast, 2C extraction reduced the overall IP signal, but a shifted band became clearly visible above the size of native GAPDH in the sample not treated with RNase. This shifted band is RNase-sensitive and the signal accumulates at the expected size of GAPDH following RNA digestion (Suppl. Figure 2). These results clearly show that 2C strongly enriches for the RNA-bound forms of GAPDH for immunoprecipitation, potentially improving the signal to noise ratio of subsequent sequencing experiments.

**Figure 2.**
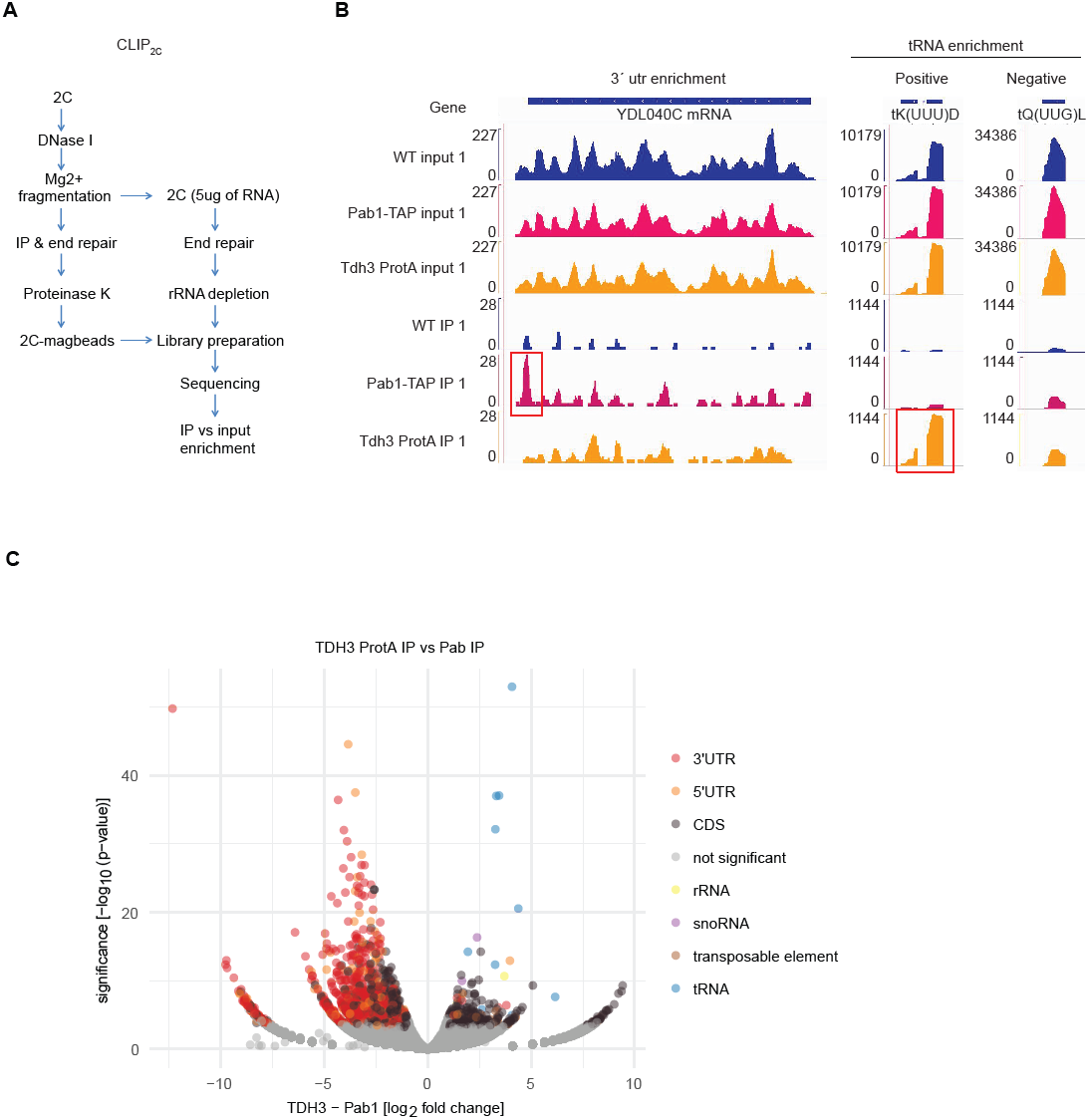
CLIP_2C_ identifies several tRNAs as specific GAPDH biding partners. **A)** Schematic representation of CLIP_2C_ workflow. **B)** Read coverage along YDL040C, tK(UUU)D and tQ(UUG)L genes after CLIP_2C_ of Pab1-TAP and Tdh3-Protein A. An untagged WT strain was used as negative control. Red boxes highlight statistically significant RNA target regions. **C)** Volcano plot displaying the log2 Fold change in normalized read counts vs -log10 p-value after CLIP_2C_ of Tdh3-Protein A and Pab1-TAP. Statistical significance was defined as Log2FC≥1 and p-value≤0.05.

Encouraged by this result, we developed CLIP_2C_ to identify the RNAs bound to GAPDH. Following a first round of 2C, the eluates were DNase I-treated and RNA was fragmented. A small aliquot was saved for sequencing an input sample, while the rest was used for immunoprecipitation. Libraries from the input and the RNA isolated from the immunoprecipitation were generated and sequenced (Figure 2A). To test the 2C-CLIP method, we used Pab1-TAP and Tdh3-Protein A tagged strains to be used in IgG-based pulldowns. An untagged WT strain was also included in the experiment as a negative control (Suppl. Figure 3). Sequencing data from the input samples were used to assess variability in gene expression between the different strains, and sequencing data from the IPs served to detect the GAPDH target RNAs. 1290 genes were identified as targets of Pab1 and as expected from a poly(A)-binding RBP, 3’utr regions were found to be particularly enriched. 326 genes, including 299 protein coding mRNAs, were identified as targets of Tdh3. This includes several tRNAs and a subset of mRNAs encoding for proteins of the glycolytic pathway (Figure 2B & table 2). However, the defined shift of RNA-crosslinked GAPDH by approximately 25 kDa suggest that the tRNAs might be preferential targets of GAPDH (Figure 1F, Suppl. Figures 2 & 3). No enrichment was observed for the WT negative control sample, strongly supporting the specificity of the results. Overall, 12 different tRNAs are significantly enriched in the Tdh3 IPs, while no tRNA was enriched in the IPs of Pab1 (Figure 2C, Table 2). Thus, similarly to the human protein, yeast GAPDH also binds tRNAs in vivo.

**Figure 3.**
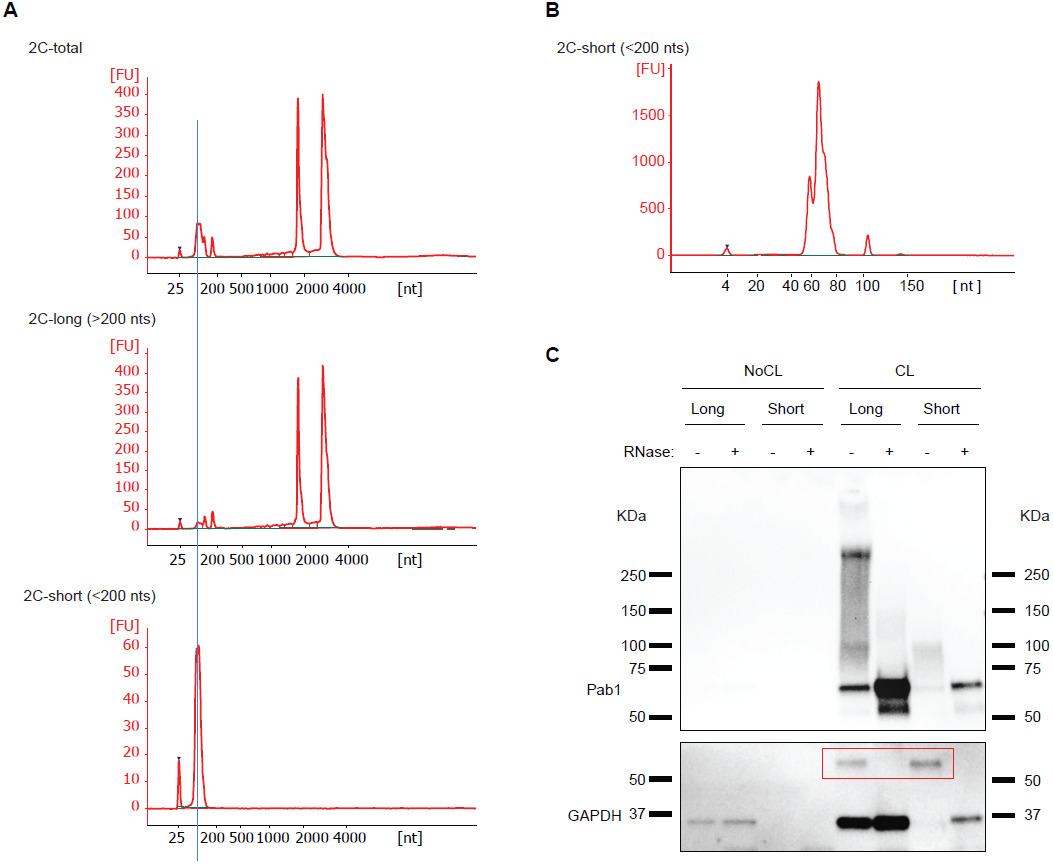
GAPDH is enriched in a fraction of short 2C-RNAs. **A)** Partition of 2C RNA based on RNA length. 2C RNA from a first extraction was subjected to a second round of 2C under conditions that retain total RNA (upper bioanalyzer electropherogram), RNA molecules longer than 200 nt (middle electropherogram) or RNA molecules shorter than 200 nt (lower electropherogram). **B)** Bioanalyzer electropherogram of 2C short RNA loaded a short RNA chip. **C)** 2C RNA from the long and short fractions from CL and NoCL samples were assayed by 2C-WB and probed against Pab1 and GAPDH antibodies. RNA loaded for the long and short fractions was 15 and 1.4ug respectively. GAPDH-RNA complexes are highlighted in the red box.

### snRIC_2C_ yields the yeast proteome of small non-coding RNA-binding proteins

Which other RBPs might preferentially bind to tRNAs or other small non-coding RNAs? We wondered whether 2C could be adapted to address this question, because the binding of small versus longer RNAs to silica matrices is known to be sensitive to the concentration of ethanol in the buffer (Hu et al.2020). Starting with a 2C total RNA eluate (Figure 3A upper panel), we separated RNAs longer (Figure 3A, middle panel) and shorter (Figure 3A, lower panel) than 200 nucleotides, respectively, during a second round of differential 2C (see methods section for experimental details). Using a bioanalyzer chip optimized to resolve small RNAs, the shorter RNA fraction is found to peak at 66 nucleotides, close to the length of yeast tRNAs (Figure 3B). While the small RNA fraction shows little if any contamination by longer RNAs (Figure 3A, bottom panel), the long RNA fraction still includes noticeable amounts of small RNAs (Figure 3A, middle panel). When 10x long (15 ug) and 1x short (1,4 ug) RNA were compared by western blotting for GAPDH following 2C, the signal of shifted GAPDH-RNA complexes from short RNA exceeds that from long RNA, reflecting a strong enrichment of crosslinked GAPDH in the 2C short RNA fraction (Figure 3C). As expected, crosslinked Pab1 is strongly enriched in the long RNA fraction, supporting the specificity of the size separation process.

Based on the successful determination of the total RNA-binding proteome by RIC_2C_ and the excellent separation between small and long RNAs using 2C, we felt encouraged to apply 2C for the determination of the first proteome-wide dataset of proteins that bind to small non-coding RNAs by snRIC_2C_ (Figure 4A). In essence, UV-crosslinked and non-crosslinked samples were subjected to a first round of 2C and eluates were treated with DNase I, as in RIC_2C_. Each sample was then split into two equal aliquots for a second round of 2C. One aliquot followed the RIC_2C_ protocol and underwent a second round of 2C for elution of total RNA, the second aliquot was used for differential 2C to isolate the small RNA fraction. Crosslinked proteins co-purified with the 2C total and small RNA fractions from the second round were then furnished with tandem mass tags (TMT) and analyzed by mass spectrometry. The raw TMT signals showed high reproducibility between experimental repeats and a clear enrichment of the total and small RNA-crosslinked samples over their respective non-crosslink controls (Suppl. Figures 4A & 4B).

**Figure 4.**
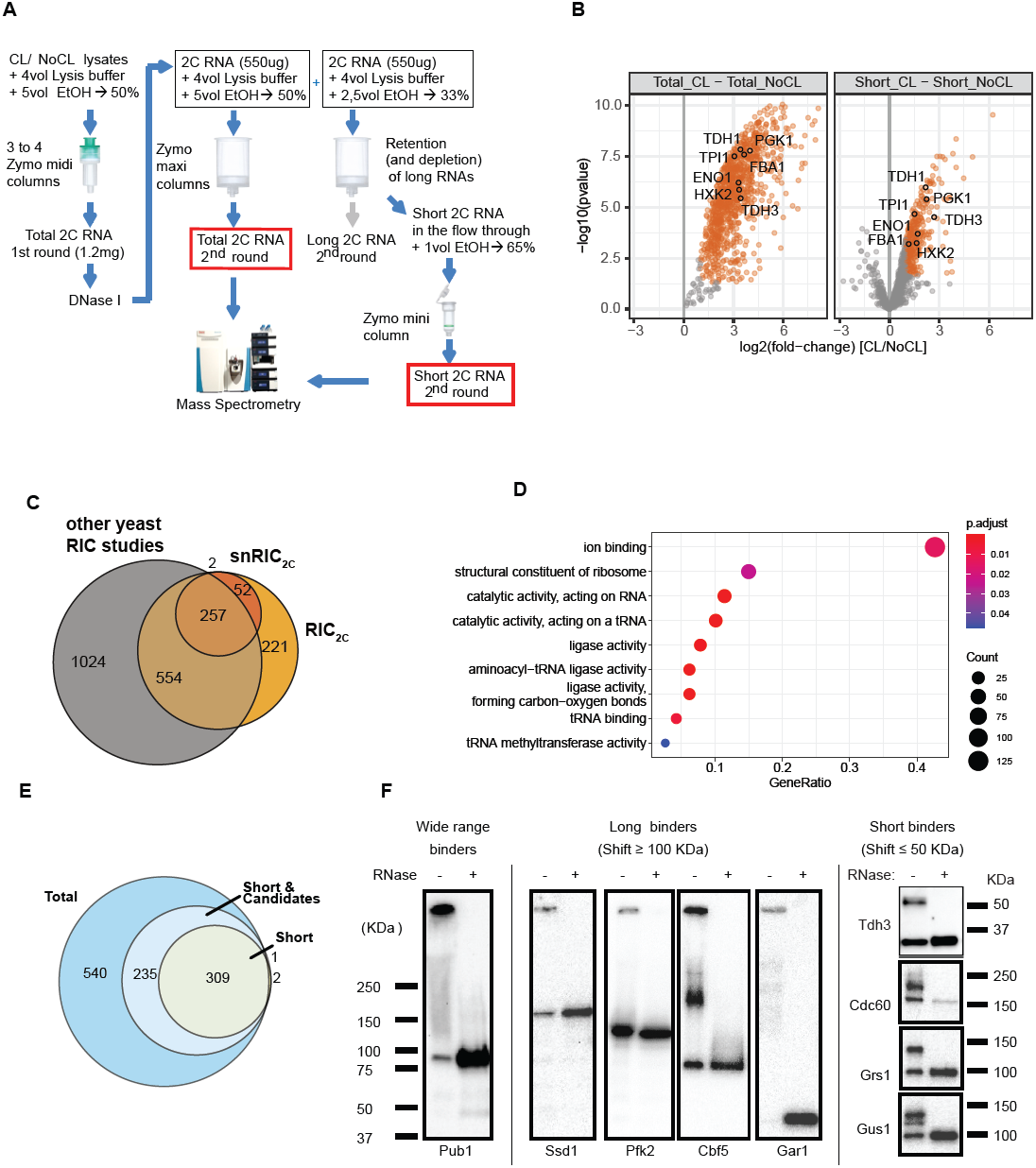
snRIC_2C_ identified 311 short RNA binding proteins in yeasts. **A)** Schematic representation of snRIC_2C_ workflow. **B)** Volcano plot displaying log2 fold change of protein abundance vs –log10 p-value for total (left panel) and short (right panel) 2C-RNA respectively. Orange dots represent statistically significantly enriched proteins in CL vs NoCL samples. Glycolytic enzymes statistically enriched in the CL RNA fractions are highlighted. **C)** Comparison of RBPs detected in the short RNA fraction vs the 2C total RNA fraction or any other yeast RIC experiment. **D)** GO term analysis of short RNA binding proteins identified by snRIC_2C_. **E)** Venn diagram classifying the RBPs identified in the snRIC_2C_ experiment. Proteins statistically enriched (logFC higher than 1 and p-value lower than 0.05) in the total RNA fraction and not enriched in the short RNA one, were considered long RNA binders. Proteins statistically enriched in the short RNA fraction were considered short RNA binders. Proteins statistically enriched in the total RNA fraction and candidate hits in the short RNA fraction (logFC between 0.5 and 1 and p-value lower than 0.05) were considered wide range RNA binders (light blue circle). **F)** Validation of the classification of RBPs based on the length of RNA target molecules by 2C-WB. TAP-tagged and an untagged WT strain were UV crosslinked and 1mg of lysate was subjected to a 2C extraction for total RNA. TAP-tagged strains were probed with a PAP antibody for Protein A detection. Untagged WT strain was probed with a GAPDH antibody. Pub1-TAP was tested as a wide range RBP. Ssd1–TAP, Pfk2-TAP, Cbf5-TAP and Gar1-TAP were tested as long RNA binders. GAPDH, in the untagged WT strain, Cdc60-TAP, Grs1-TAP and Gus1-TAP were tested as short RNA binders.

Around 1000 RBPs were identified in the total RNA samples with an overlap of over 75% with the original RIC_2C_ experiment (Suppl. Figure 4C), reflecting the reproducibility of the method. snRIC_2C_ identified 311 proteins that bind to purified small RNAs (Figure 4B and C, table 3), including 52 RBPs that were not previously annotated as such in yeast (Figure 4C). Subsequent GO-term analysis of these proteins showed a strong enrichment for terms related to tRNA metabolism (Figure 4D), as expected, supporting the validity of snRIC_2C_.

While the differential 2C elution yields quite pure small RNA fractions without noticeable long RNA contamination, the converse does not apply to the long RNA fraction (Figure 3A). Therefore, we only considered the crosslinked proteomes associated with total and small RNA for comparative analyses (Figure 4E). Together with the 311 highly enriched proteins in the short RNA fraction, we found 235 additional proteins that fell slightly below the statistical threshold for significance in snRIC_2C_ (log2FC≥1 and p-value≤0.05) but were strongly enriched in RIC_2C_. We suggest that these RBPs likely bind long and small RNAs (‘wide range RBPs’). Finally, 540 RBPs were detected by RIC_2C_ but undetected in snRIC_2C_ samples and, therefore, are considered as preferential long RNA binders (Figure 4E).

To validate these data, we used different TAP-tagged strains and examined UV-crosslinked and non-crosslinked samples of these by Western blot following 2C. In addition to GAPDH (Tdh3), three tRNA synthetases identified as small RNA binders were examined: Cdc60, Grs1 and Gus1. As observed before with GAPDH, all four RBPs show a sharp, RNase-sensitive additional band migrating <50 kDa more slowly than the native proteins before RNase treatment (Figure 4F). By contrast, the ‘wide range binder’ Pub1 and the long RNA binders Ssd1, Pfk2, Cbf5 and Gar1 all show different patterns with RNase-sensitive smears and/or additional bands migrating >100 kDa more slowly or being retained in the wells of the gel. These results support the assignment of small non-coding RNA binders by snRIC_2C_, and confirm that the shifts observed in 2C immunoblots correlate well with the lengths and homo-/heterogeneity of the bound RNAs.

### Analysis of small non-coding RBPs reveals enrichment of glycolytic and TCA cycle enzymes

Classification of the snRBPs shows that 41% of the 311 RBPs are enzymes, including 45 metabolic enzymes (Figure 5A). Amongst the metabolic enzymes, processes related to tRNA metabolism and carbohydrate derivative metabolic processes are strongly enriched (Figure 5B). Strikingly, 10 glycolytic enzymes (Hxk2, Fba1, Tdh1-3, Tpi1, Pgk1, Gpm1 and Eno1-2) and five enzymes of the TCA cycle (Aco1, Idh1, Mdh1, Lsc1 and Lpd1) were identified as snRBPs (Figure 4B & 5C), and aminoacyl-tRNA biosynthesis and carbon metabolism are the two significantly enriched pathways amongst the snRIC_2C_ hits (Figure 5D). To confirm this striking enrichment, we tested several further TAP-tagged strains in 2C western blot experiments, using an untagged WT strain as a negative control (Suppl. Figure 5). These experiments confirm that the glycolytic enzymes Tdh3, Fba1, Pgk1, Hxk2, Gpm1, Tpi1 and Eno1 all show the defined, RNase-sensitive additional band less than 50 KDa larger than the expected size of the respective proteins (Figure 5E). We conclude that several yeast glycolytic enzymes bind small non-coding RNAs in addition to their well-known roles in central carbon metabolism.

**Figure 5.**
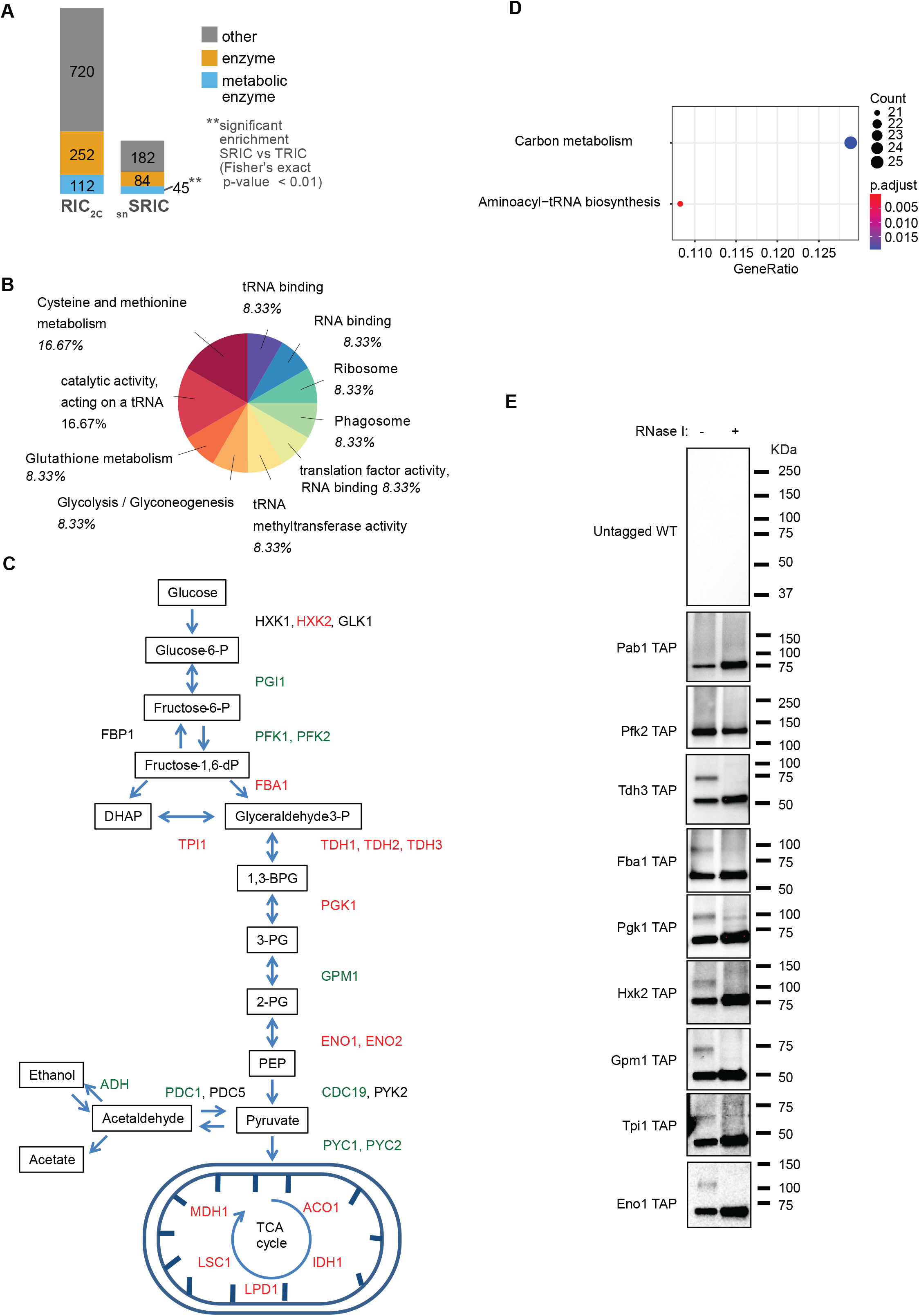
Glycolytic pathway is enriched in short RNA binding proteins. **A)** Analysis of proteins found in snRIC_2C_ experiment. **B)** Go-term molecular function and interpro domain analysis of the proteins enriched in the CL short fraction in snRIC_2C_ experiment. **C)** Schematic representation of glycolysis and TCA cycle pathways. Proteins highlighted in green were identified as long or wide range RBPs and proteins highlighted in red were found to bind short RNAs in snRIC_2C_. **D)** Analysis of KEGG enriched pathways after snRIC_2C_. Aminoacyl-tRNA biosynthesis and carbon metabolism pathways were found to be statistically enriched from the proteins detected in the short RNA fraction in snRIC_2C_ experiment. **E)** Validation of glycolytic and TCA enzymes as short RBPs by 2C-WB. TAP-tagged strains and an untagged WT strain were UV-crosslinked and 1mg of lysate was used in a round of total 2C-RNA extraction. Blots were probed against PAP antibody. The untagged WT strain was tested as a negative control for the western blot. Pab1-TAP and Pfk2-TAP were used as controls as wide range and long RNA binders respectively. Tdh3-TAP, Fba1-TAP, Pgk1-TAP, Hxk2-TAP, Gpm1-TAP, Tpi1-TAP and Eno1-TAP were tested as short RNA binders.

### Small non-coding RNA binding of GAPDH is carbon source-dependent and regulated by Maf1

We wondered whether the binding of glycolytic enzymes to small non-coding RNAs is constitutive or subject to biological regulation. A growing body of evidence links nutrient availability with the regulation of RNA polymerase III activity by its universal repressor Maf1 (Graczyk et al.2018, Morawiec et al.2013, Willis2018). Under favorable growth conditions like fermentation, Maf1 is repressed, allowing high Pol III activity and tRNA transcription. Conversely, growth under respiratory conditions results in Maf1 activation, which represses Pol III activity and tRNA synthesis (Ciesla et al.2007, Graczyk et al.2018, Morawiec et al.2013, Willis2018). Therefore, we tested whether the tRNA binding activity of GAPDH responds to changes of the carbon source in the growth media and is affected by Maf1. When WT and Maf1 KO cells were grown in glucose, a fermentative carbon source, or glycerol and ethanol, which can only be respired, we observed tRNA engagement of GAPDH only under fermentative conditions (Figure 6A). Under these conditions, RNA binding is unaffected by Maf1 deletion. However, deletion of Maf1 strongly induces the binding of GAPDH to tRNA under respiratory conditions, effectively suppressing its regulation by carbon source. These results uncover a connection between tRNA binding of glycolytic enzymes, the activity of the glycolytic pathway that is influenced by the prevalent carbon source and Pol III activity (Figure 6B).

**Figure 6.**
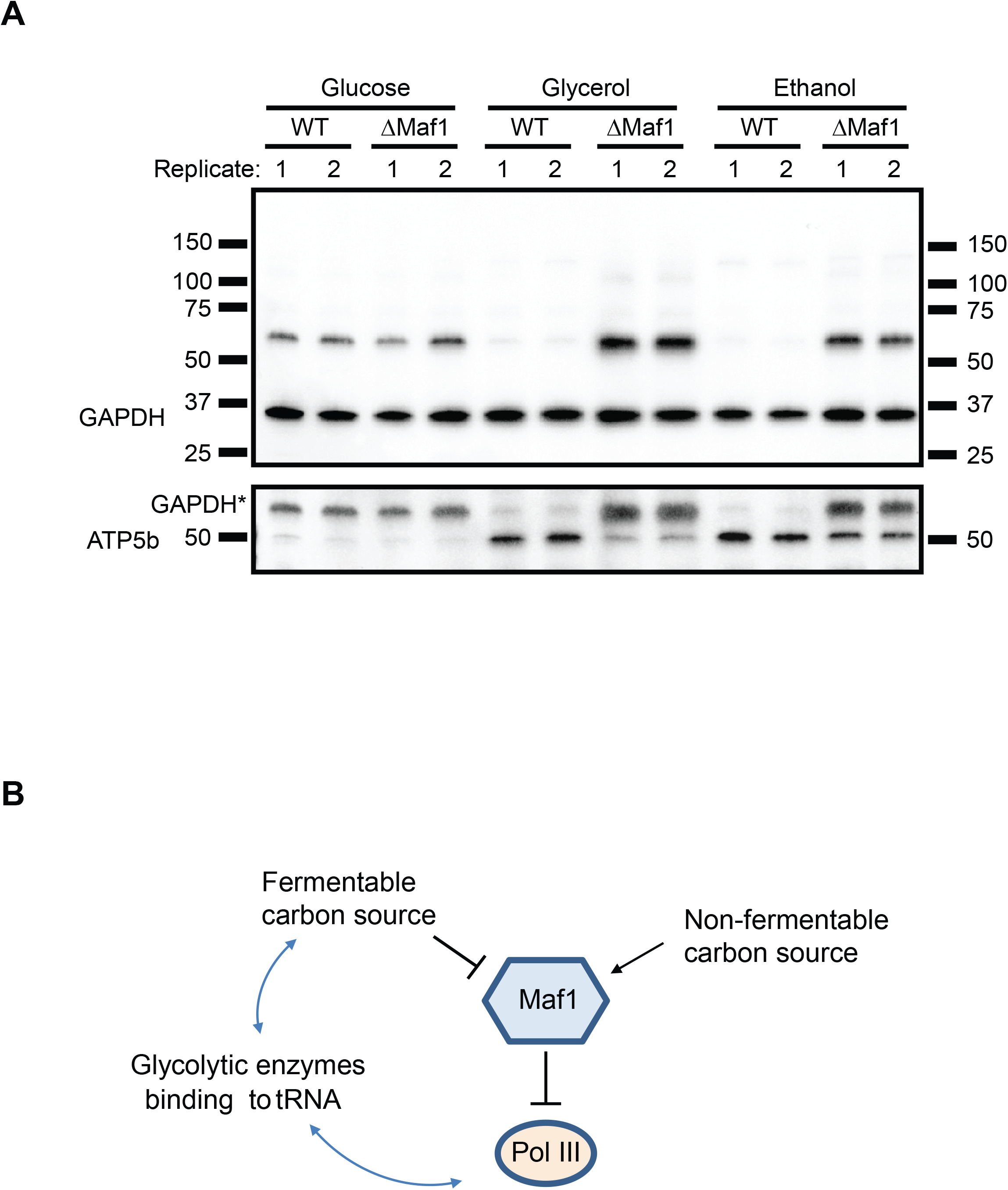
Engagement of GAPDH to tRNAs is carbon source-dependent and regulated by the RNA polymerase III universal repressor Maf1. **A)** WT and ΔMaf1 strains were grown in glucose, glycerol and ethanol and UV-crosslinked. Total 2C RNA extractions were performed from two different biological replicates and 10ug of RNAs were tested on 2C-WBs against GAPDH and ATP5b antibodies. GAPDH* represents residual GAPDH signal after stripping and reprobing the membrane. **B)** The RNA binding activity of glycolytic enzymes link the nutrient-dependent regulation of Maf-1 with the products of RNA PolIII. Maf1, universal repressor of RNA polymerase III activity, is regulated by different metabolic signals. Maf1 is activated in response to DNA damage and is inhibited in the presence of oncogenic transformation, growth factors and nutrients. Glycolytic enzymes bind both, nutrients and the products of PolIII activity in the form of tRNAs and might represent a regulatory link of both processes.

## Discussion

We previously showed that commercially available silica columns for the purification for total cellular RNA can also be used to select for RBPs that are covalently crosslinked to these RNAs, offering a simple method (called complex capture [2C]) to test the RNA-binding activity of proteins by simple immunoblotting of 2C eluates (Asencio et al., 2018). Realizing that the 2C principle can also be used to determine the total RNA-binding proteomes of cells if eluates are analyzed by sensitive mass spectrometry, we applied this method to the yeast *Saccharomyces cerevisiae*. Our unexpected biological observations drove further methodological advances, and this work hence reports both the development of enabling methods and new biological insights on the interaction of glycolytic enzymes with small non-coding RNAs in yeast.

### Methodological advances

*RIC*_*2C*_. Based on our earlier work, 2C could be readily applied to the determination of the proteome that binds to any class of cellular RNAs. For yeast, we found a total number of 983 RBPs, which is in keeping with other reports both in terms of the number and the identity of the RBPs. This result both validates RIC_2C_ methodologically and suggests that at least under standard growth conditions the number of yeast RBPs appears to approximate saturation, since we identified only a modest number of 174 RBPs that were not detected previously. RIC_2C_ provides advantages compared to alternative methods. It does not require in vivo labelling of RNA with nucleotide analogues, like CARIC (Huang et al.2018). RIC_2C_ is simple, easily scalable and does not require the challenging isolation of RBPs from the interphase between two solvents, as in OOPS, XRNAX and PTex (Queiroz et al.2019, Trendel et al.2019, Urdaneta et al.2019). Moreover, organic phase separation methods like OOPS can under-represent RBPs bound to small RNAs (Queiroz et al.2019). RIC_2C_ conceptually corresponds to the recently published TRAPP protocol (Shchepachev et al.2019), where silica powder is used as the starting material for the enrichment of total RNA-binding proteins. RIC_2C_ does not require extensive washing and preparation steps for the purification columns prior to the application of samples, because it uses columns and buffers contained in commercially available RNA extraction kits. Therefore, it is a simple and straightforward method that can be applied to a wide range of biological materials, both eukaryotic and prokaryotic in origin. Because different column sizes are commercially available, it is simple to scale RIC_2C_ to the experimental needs.

*CLIP*_*2C*_. UV-<u>c</u>rosslinking followed by <u>i</u>mmuno<u>p</u>recipitation and library preparation from the co-precipitated RNAs is commonly used to determine the RNAs bound to an RBP of interest (Darnell2010, Lee & Ule2018, Ule et al.2018, Van Nostrand et al.2016). While CLIP protocols typically perform well when studying canonical, high affinity RBPs such as e.g. RNA processing factors, they can fall short with non-canonical, lower affinity RBPs, where often only a minor fraction of the protein is RNA-bound which can give rise to a high non-specific background. CLIP_2C_ offers a simple enrichment step for the RNA-bound fraction of an RBP of interest (Figure 2A & S2) before immunoprecipitation. The resulting RNA-loaded RBP subsequently represents an ideal substrate for library generation and sequencing, because contaminant proteins in the immunoprecipitation are reduced. Nonetheless, 2C requires protein denaturing conditions, which potentially compromises the subsequent immunoprecipitation step if the antibody used recognizes a natively folded epitope. Therefore, antigen-antibody pairs should be pre-evaluated for their compatibility with denatured/renatured proteins. We also show that proteins bearing protein-A tags can be efficiently pulled-down from 2C eluates, subsequently yielding excellent sequencing results. We used CLIP_2C_ to demonstrate for the first time that yeast GAPDH binds tRNAs in vivo, supporting the evolutionary conservation of the GAPDH-tRNA interaction previously reported for human GAPDH (Singh & Green1993). *snRIC*_*2C*_. Unlike other methods for the capture of RNA-binding proteomes, the silica matrix-based approach allows robust and reproducible separation of small (<200 nucleotides) from longer RNAs. We show here that based on this principle snRIC_2C_ can be used to determine the collective of RBPs that binds to small non-coding RNAs. As discussed below, this collective displays interesting distinctions from the total RNA-bound proteome as a whole. To the best of our knowledge, snRIC_2C_ represents the first method for the systematic isolation of RBPs based on the lengths of their target RNAs. snRIC_2C_ may also be of particular interest for the field of bacterial small RNA metabolism, as small non-coding RNAs are highly recognized for their critical regulatory roles in bacteria (Jørgensen et al.2020, Ponath et al.2022, Quendera et al.2020), but methods to identify and study their associated RBPs are still needed.

### New biological datasets and insights

#### Identification of the snRBPs from yeast

Using snRIC_2C_, we identified ∼300 yeast proteins that are highly enriched for binding to small non-coding RNAs, snRBPs (Figure 5). Since small non-coding RNAs exert numerous regulatory functions, it is important to reveal the snRBPs with which they preferentially interact. Unsurprisingly, the snRBPs include many proteins known for their roles in tRNA metabolism and function. But it is quite unexpected to find so many glycolytic and TCA cycle enzymes amongst the snRBPs (Figure 4B & 5C). Earlier work connected yeast glycolytic enzymes with tRNAs. For instance, enolase has been found to bind tRNAs in vivo (Shchepachev et al.2019) and has been proposed to participate in the import of tRNA to mitochondria in yeast (Entelis et al.2006). Interestingly, while enolase was originally considered to directly contribute to the import of tRNA into mitochondria, later publications favour the view that enolase accompanies other proteins participating in this process (Baleva et al.2017). Our results broadly implicate glycolytic enzymes and TCA cycle enzymes in the binding of small non-coding RNAs. At least for GAPDH, these largely appear to be tRNAs (Figure 2B, 2C). These observations raise the question of what the function(s) of these RNA-protein interactions may be. As exemplified above and by other examples, the enzymes may moonlight in critical aspects of small non-coding RNA biology. Alternatively, the RNAs may riboregulate the enzymes that they bind to. Riboregulation has recently been shown for human enolase 1 (Huppertz et al.2022), and the human small non-coding vtRNA1-1 has been identified to regulate mammalian autophagy by binding to the receptor protein p62 (Horos et al., 2019).

With ∼300 yeast snRBPs having been identified and the application of snRIC_2C_ to other organisms, we expect further insights into the biological functions of small non-coding RNAs.

#### A carbon source-regulated interaction between glycolysis and RNA polymerase III activity

Cells dedicate profound resources to protein production, not only involving translation itself, but also including the transcription, maturation and amino-acylation of tRNAs. Therefore, cells must monitor nutrient availability and modulate protein and tRNA synthesis accordingly. The activity of the tRNA synthesizing Pol III is inhibited under nutrient limiting conditions by the repressor protein Maf1 (Upadhya et al.2002). Here, we show that the binding of tRNAs to GAPDH is regulated by Maf1. Earlier work identified regulatory interactions between carbon metabolism enzymes and Pol III activity (Ciesla et al.2007, Morawiec et al.2013, Szatkowska et al.2019), but fell short of noticing the direct and carbon source-dependent interaction of glycolytic enzymes with Pol III transcripts. Our findings may indicate a critical link between nutrient availability, energy metabolism and Pol III activity (Figure 6B). More research will serve to analyze these interactions in detail.

## Supporting information

Supp table 1_compendium of RIC datasets

Table 1. RIC2C Limma results

Table 2. CLIP2C_target genes

Table 3_snRIC2C_Limma results

## Acknowledgements

We thank Dr. T. Sekaran for his support with the data analysis, and current and former members of the Hentze laboratory for their feedback, suggestions and fruitful discussions. We thank the EMBL Genomics, Proteomics and Protein Expression and Purification core facilities for their expert support. We thank C. Girardot for helping with sequencing data deposition. We are greatful to Dr. J. Heinisch for the provision of the anti-Pfk antibody. This work was supported in part by the Manfred Lautenschläger Foundation (to MWH).

## Figure legends

**Supplementary figure 1.**
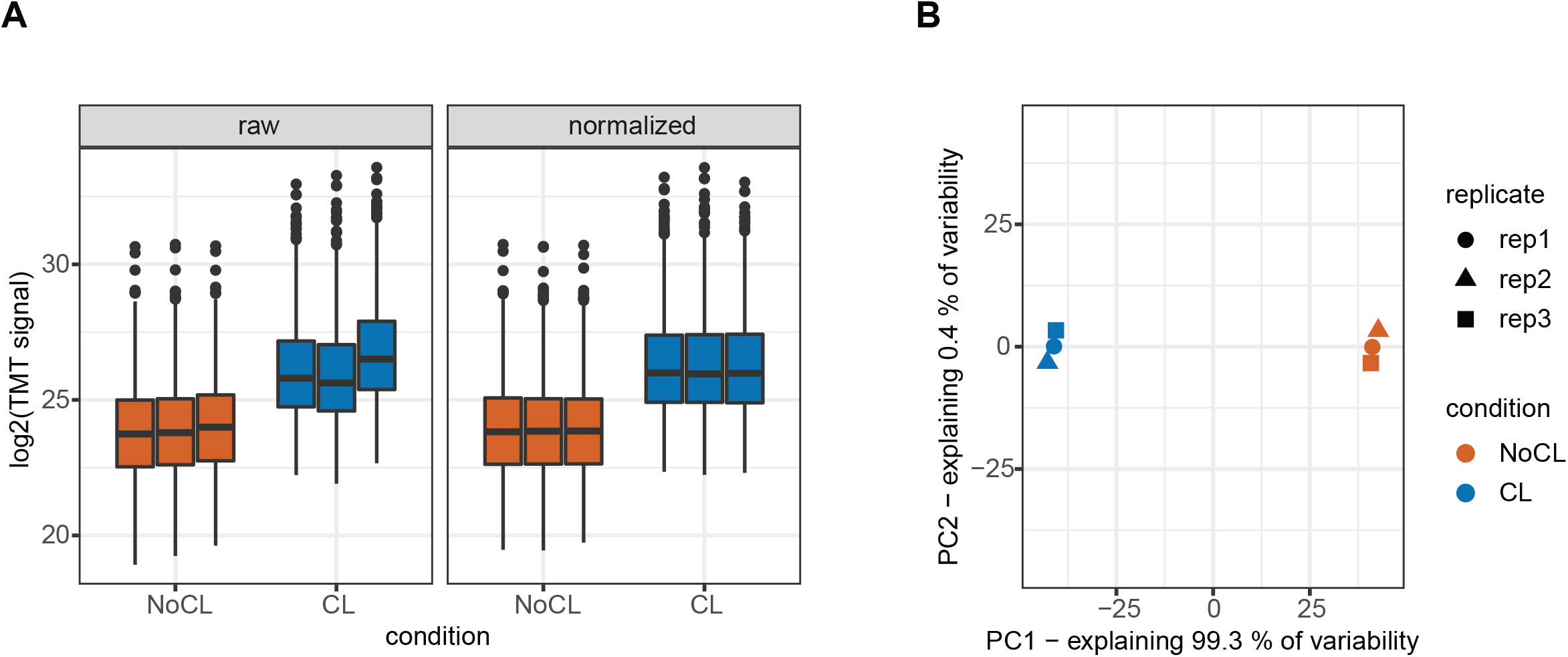
Quality control of RIC_2C_. TMT raw and normalized signal sum **A)** and principal component analysis **B)** of three biological replicates for CL and NoCL samples.

**Supplementary figure 2.**
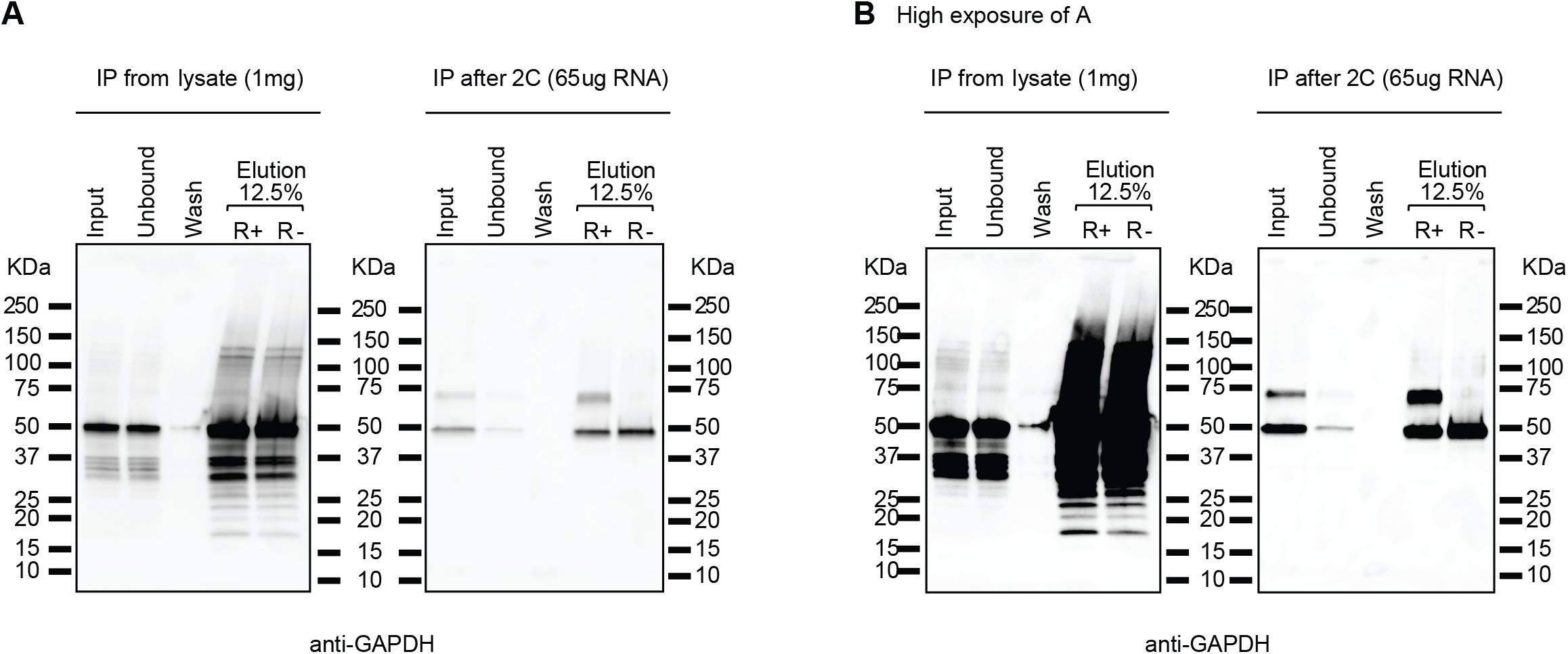
Comparison of IP efficiency from a lysate or from a 2C eluate. Pull downs of Tdh3-Protein A were done from 1mg of protein lysate and 65 ug of 2C RNA obtained from crosslinked samples and non-crosslinked negative controls. Input, unbound, wash and eluate fractions were separated by SDS-PAGE and probed with an anti-GAPDH antibody. **A)** Low exposure. **B)** High exposure.

**Supplementary figure 3.**
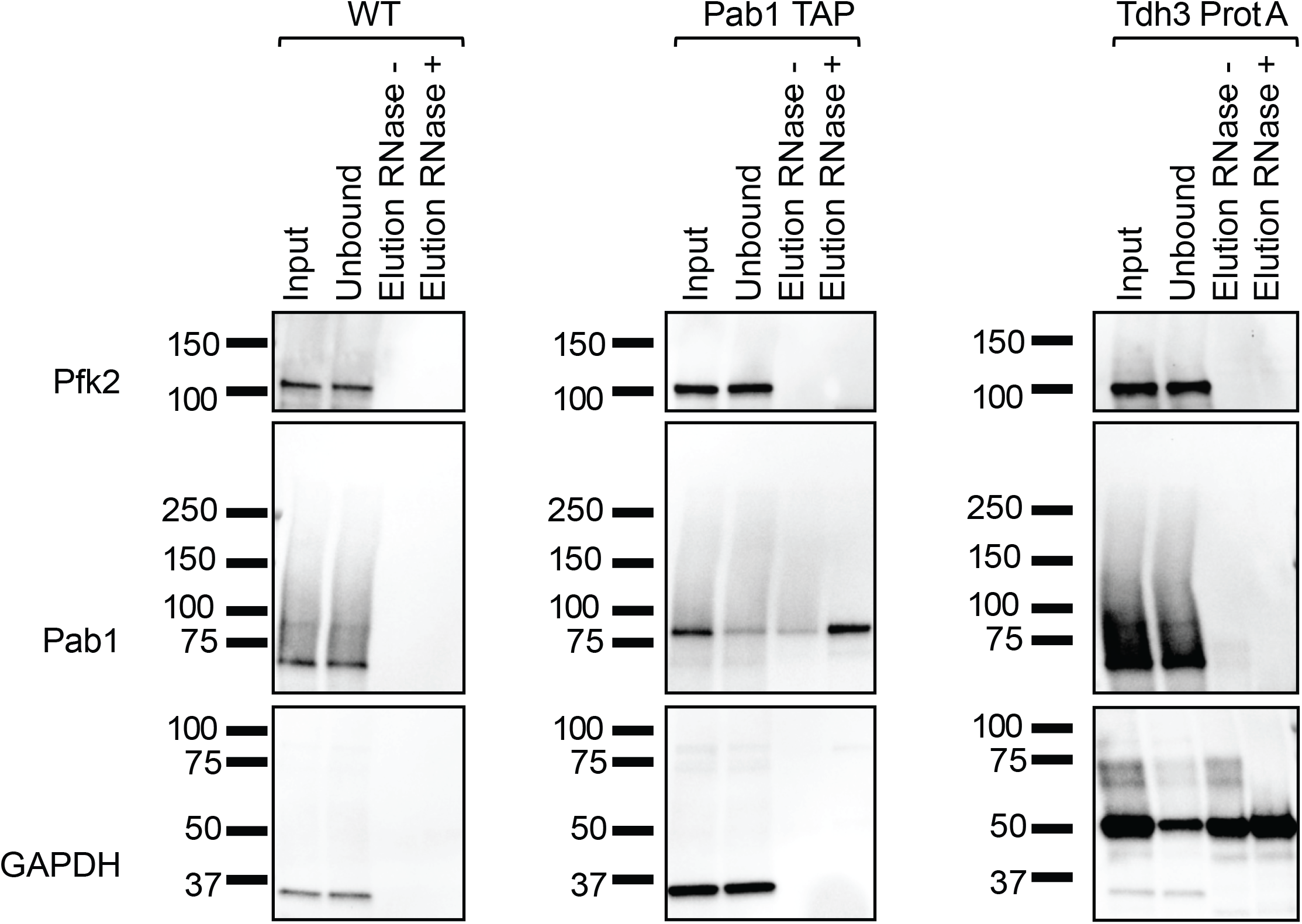
Validation of IPs from CLIP_2C_ experiment. Performance of the CLIP_2C_ pulldowns for WT, Pab1-TAP and Tdh3-Protein A was assessed by western blot. Aliquots of input, unbound and elution fractions were separated by SDS-PAGE and probed against Pfk, Pab1 and GAPDH antibodies.

**Supplementary figure 4.**
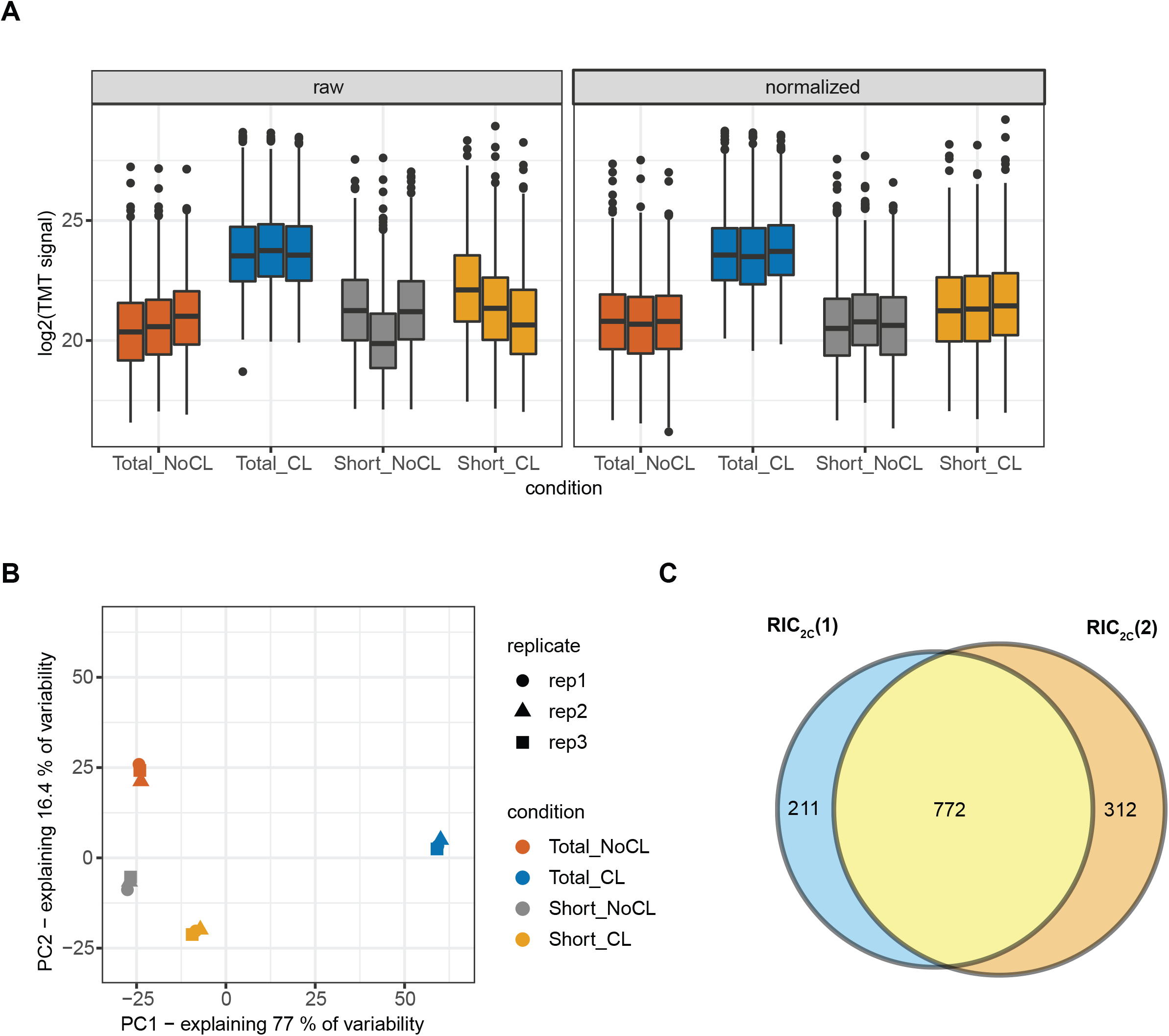
Quality control of snRIC_2C_. TMT raw and normalized signal sum **A)** and principal component analysis **B)** of total and short RNA for CL and NoCL samples. **C)** Venn diagram comparing original RIC_2C_ experiment (RIC_2C_(1), see also Figure 1) with the proteins detected in 2C total RNA fraction (RIC_2C_(2)) from the snRIC_2C_ experiment.

**Supplementary figure 5.**
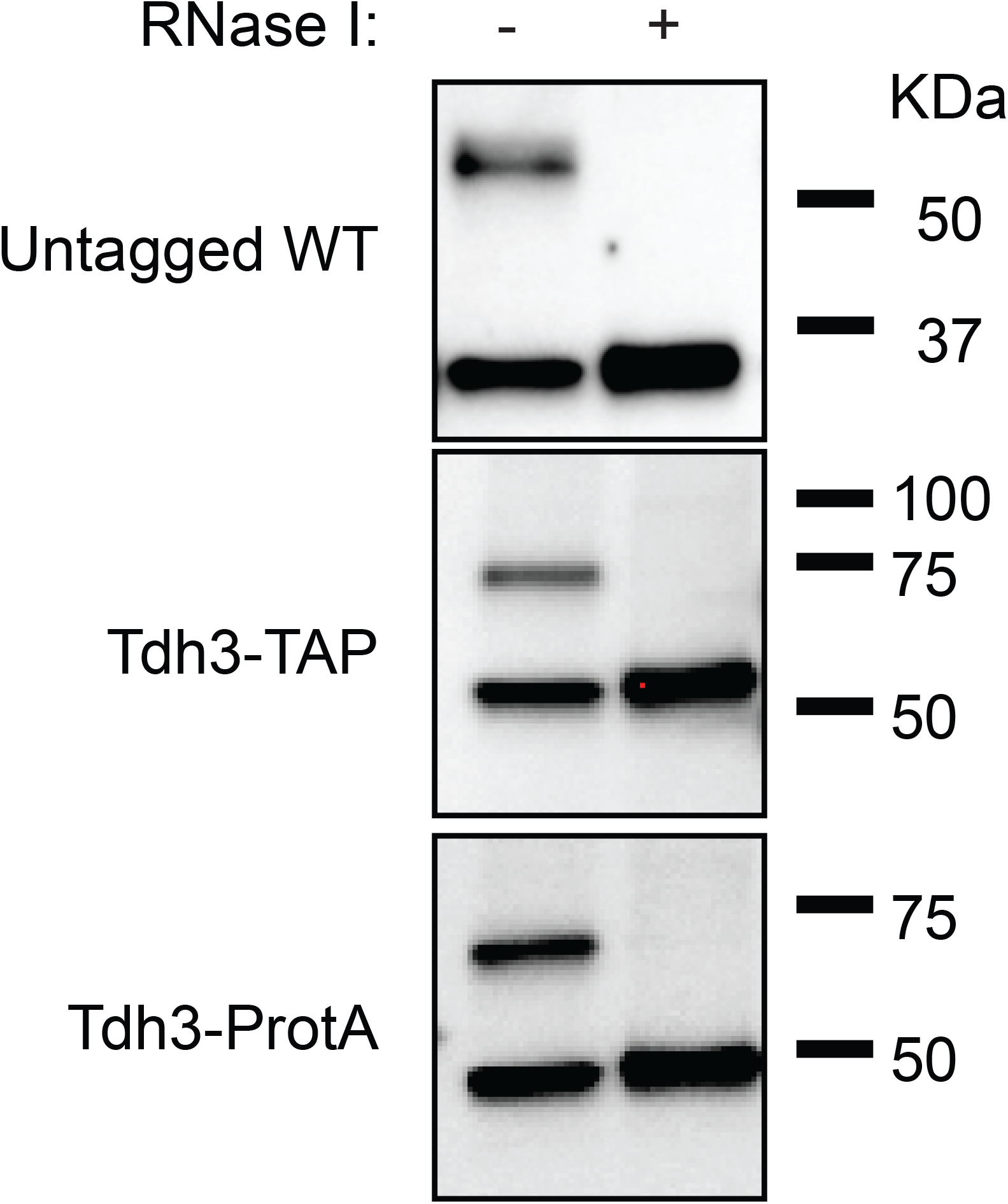
Validation of short RNA binding of GAPDH in an untagged WT strain and in Tdh3-TAP and Tdh3-Protein A tagged strains. An untagged WT strain, a Tdh3-TAP and a Tdh3-Protein A tagged strains were UV-crosslinked to test the effect of different tags on the binding of GAPDH to RNA. 1mg of protein lysate was used on a total 2C-RNA extraction and 10ug of RNA were treated or not with RNase I and tested on a 2C-WB experiment. The untagged WT strain was probed against a GAPDH antibody. Tdh3-TAP and Tdh3-Protein A strains were probed against PAP antibody.

## Methods

### *S. cerevisiae* strains and manipulations

Standard methods were used for yeast culture and manipulation (Amberg et al.2005). Yeast strains, genotype and origin are summarized in the following table:

| Strain | Genotype | Source |
| --- | --- | --- |
| WT BY4741 | MATa his3Δ1 leu2Δ0 met15Δ0 ura3Δ0 | Horizon<br>Discovery<br>LTD |
| ΔMaf1 | MATa his3Δ1 leu2Δ0 met15Δ0 ura3Δ0 maf1Δ | Horizon<br>Discovery<br>LTD |
| Cbf5-TAP | MATa his3Δ1 leu2Δ0 met15Δ0 ura3Δ0 CBF5-TAP::His3Mx6 | Horizon<br>Discovery<br>LTD |
| Cdc60-TAP | MATa his3Δ1 leu2Δ0 met15Δ0 ura3Δ0 CDC60-TAP::His3Mx6 | Horizon<br>Discovery<br>LTD |
| Eno1-TAP | MATa his3Δ1 leu2Δ0 met15Δ0 ura3Δ0 ENO1-TAP::His3Mx6 | Horizon<br>Discovery<br>LTD |
| Fba1-TAP | MATa his3Δ1 leu2Δ0 met15Δ0 ura3Δ0 FBA1-TAP::His3Mx6 | Horizon<br>Discovery<br>LTD |
| Gar1-TAP | MATa his3Δ1 leu2Δ0 met15Δ0 ura3Δ0 GAR1-TAP::His3Mx6 | Horizon<br>Discovery<br>LTD |
| Gpm1-TAP | MATa his3Δ1 leu2Δ0 met15Δ0 ura3Δ0 GPM1-TAP::His3Mx6 | Horizon<br>Discovery<br>LTD |
| Grs1-TAP | MATa his3Δ1 leu2Δ0 met15Δ0 ura3Δ0 GRS1-TAP::His3Mx6 | Horizon<br>Discovery<br>LTD |
| Gus1-TAP | MATa his3Δ1 leu2Δ0 met15Δ0 ura3Δ0 GUS1-TAP::His3Mx6 | Horizon<br>Discovery<br>LTD |
| Hxk2-TAP | MATa his3Δ1 leu2Δ0 met15Δ0 ura3Δ0 HXK2-TAP::His3Mx6 | Horizon<br>Discovery<br>LTD |
| Pab1-TAP | MATa his3Δ1 leu2Δ0 met15Δ0 ura3Δ0 PAB1-TAP::His3Mx6 | Horizon<br>Discovery<br>LTD |
| Pfk2-TAP | MATa his3Δ1 leu2Δ0 met15Δ0 ura3Δ0 PFK2-TAP::His3Mx6 | Horizon<br>Discovery<br>LTD |
| Pgk1-TAP | MATa his3Δ1 leu2Δ0 met15Δ0 ura3Δ0 PGK1-TAP::His3Mx6 | Horizon<br>Discovery<br>LTD |
| Pub1-TAP | MATa his3Δ1 leu2Δ0 met15Δ0 ura3Δ0 PUB1-TAP::His3Mx6 | Horizon<br>Discovery<br>LTD |
| Ssd1-TAP | MATa his3Δ1 leu2Δ0 met15Δ0 ura3Δ0 SSD1-TAP::His3Mx6 | Horizon<br>Discovery<br>LTD |
| Tdh3<br>Protein A | MATa his3Δ1 leu2Δ0 met15Δ0 ura3Δ0 TDH3-ProtA::His3Mx6 | Asencio<br>et<br>al.2018 |
| Tdh3-TAP | MATa his3Δ1 leu2Δ0 met15Δ0 ura3Δ0 TDH3-TAP::His3Mx6 | Horizon<br>Discovery<br>LTD |
| Tpi1-TAP | MATa his3Δ1 leu2Δ0 met15Δ0 ura3Δ0 TPI1-TAP::His3Mx6 | Horizon<br>Discovery<br>LTD |

### Yeast culture, cross-linking, lysate preparation, 2C method and 2C-Western blot

Yeast culture, UV cross-linking, cell lysis, 2C method and 2C western blot experiments were done as previously described (Asencio et al.2018). The following antibodies were used in western blot experiments: Anti-Pab1 1:4000 (Abcam, #ab189635), Anti-GAPDH 1:4000 (Sigma-Aldrich #G9545, [33]), Anti-Histone H3 HRP 1:1000 (Abcam, #ab21054,) Anti-Hexokinase 1:10000 (Biorad, #4959-9988), Anti-Tpi 1:4000 (Proteintech, #10713-1-AP) and Anti-tubulin 1:4000 (Abcam, #ab6160), Anti-PFK antibody 1:4000 (Heinisch1986). Peroxidase anti-peroxidase (PAP) antibody 1:10000 (Sigma-Aldrich # P1291).

### Up and downscaling the 2C method

In our original description of the 2C method, we used Zymo-Spin V-E (#C1024; Zymo Research) columns which are included in Zymo Research RNA extraction midi kits or can be purchased separately. However, the 2C method can be up or downscaled depending on the required amount of 2C RNA and/or the available input material. We have successfully tested Zymo Resaerch micro (Zymo-Spin IC, #C1004) mini (Zymo-Spin IIICG, #C1006) and maxi (Zymo-Spin VI, #C1013) columns. Alternatively, several columns can be used in parallel to process different aliquots of the same lysate simultaneously.

### 2C total RNA interactome capture (RIC_2C_)

WT yeast cells were grown, UV-crosslinked and lysed as described above. Non-irradiated cells were cultured and processed in parallel throughout the experiment as negative controls. Three biological replicates were included in the experiment. A first round of 2C extraction was performed from 1 mg of protein lysate with Zymo-Spin V-E columns and 2C-RNA was later eluted with 300ul of nuclease-free water. Although in our hands, the DNA contamination after a 2C extraction from a yeast lysate is below 2%, we nevertheless treated the eluates with 20U of DNase I (AM2224, Ambion) at 37ºC for 30 minutes to minimize the chances of detecting DNA binding proteins after mass spectrometry analysis. A second round of 2C was performed to eliminate the DNase I enzyme and any contaminant DNA binding protein. For this, four volumes of RNA lysis buffer were added to the DNase I treated samples. Samples were mixed, five volumes of ethanol were added and after mixing, the samples were added to a second Zymo-Spin V-E column. Second round 2C-RNA was eluted with 300ul of nuclease-free water. 100ug of 2C RNA eluates were RNAse I treated and processed for TMT labelling. Briefly, cysteine’s were reduced with dithiothreitol at 56°C for 30 minutes (10 mM in 50 mM HEPES, pH 8.5) and further alkylated with 2-chloroacetamide at room temperature in the dark for another 30 minutes (20 mM in 50 mM HEPES, pH 8.5). Samples were processed using the SP3 protocol (Hughes et al.2014) and on-bead digested with trypsin (sequencing grade, Promega), which was added in an enzyme to protein ratio 1:50 for overnight digestion at 37°C. Peptides were modified with TMT6plex (Dayon et al.2008) Isobaric Label Reagent (ThermoFisher) following manufacturer’s instructions. For sample clean up, an OASIS® HLB µElution Plate (Waters) was used. Offline high pH reverse phase fractionation was performed out on an Agilent 1200 Infinity high-performance liquid chromatography system, equipped with a Gemini C18 column (3 μm, 110 Å, 100 × 1.0 mm, Phenomenex), resulting in 5 fractions.

For mass spectrometry data acquisition, an UltiMate 3000 RSLC nano LC system (Dionex) fitted with a trapping cartridge (µ-Precolumn C18 PepMap 100, 5µm, 300 µm i.d. x 5 mm, 100 Å) and an analytical column (nanoEase™ M/Z HSS T3 column 75 µm x 250 mm C18, 1.8 µm, 100 Å, Waters) was used. Trapping was carried out with a constant flow of trapping solution (0.05% trifluoroacetic acid in water) at 30 µL/min onto the trapping column for 6 minutes. Subsequently, peptides were eluted via the analytical column running solvent A (0.1% formic acid in water) with a constant flow of 0.3 µL/min, with increasing percentage of solvent B (0.1% formic acid in acetonitrile) from 2% to 4% in 4 min, then 4% to 8% in 2min, from 8% to 28% for a further 66 min, in another 10 min. from 28% to 40%, followed by an increase of B from 40-80% for 3min. and a re-equilibration back to 2% B for 5min. The outlet of the analytical column was coupled directly to an QExactive plus Mass Spectrometer (Thermo) using the Nanospray Flex™ ion source in positive ion mode.

### Mass spectrometry analysis for RIC_2C_ and snRIC_2C_ experiments

IsobarQuant (Franken et al.2015) and Mascot (v2.2.07) were chosen for data processing. A Uniprot *Saccharomyces cerevisiae* proteome database (UP000002311) containing common contaminants and reversed sequences was used. The search parameters were the following: Carbamidomethyl (C) and TMT10 (K) (fixed modification), Acetyl (N-term), Oxidation (M) and TMT10 (N-term) (variable modifications). A mass error tolerance of 10 ppm was set for the full scan (MS1) and for MS/MS (MS2) spectra of 0.02 Da. Trypsin was selected as protease with an allowance of maximum two missed cleavages. A minimum peptide length of seven amino acids and at least two unique peptides were required for individual protein identification. The false discovery rate on peptide and protein level was set to 0.01.

### CLIP_2C_

Pab1-TAP, Tdh3-Protein A and a WT untagged strain were grown in YPD, UV crosslinked and lysed as described above. A first round of 2C was performed from 1mg of protein lysate and 100ug of the obtained 2C-RNA were DNase I treated for 30 minutes at 37ºC. After DNaseI treatment, RNA samples were diluted with RNA fragmentation buffer to a final concentration of 20mM TrisHCl pH7.5, 1% SDS and 30mM MgCl2. RNA was fragmented by incubating the samples 15 minutes at 95ºC. Fragmentation was stopped by adding EDTA to a final concentration of 30mM and quickly cooling down the samples on ice and later kept at room temperature. Samples were brought to 2ml with buffer B (25mM TrisHCl 7.5mM; 140mM NaCl, 1.8mM MgCl2; 0.5mM DTT and 0.1% NP-40) and 50 and 100ul were saved for IP validation and input sequencing respectively. To the remaining volume, 100ul of pre-washed Dynabeads Pan mouse IgG (#11041, Thermo) were added per sample and were incubated at 4ºC for 2 hours with gentle rotation. Samples were washed once with buffer B, three times with wash buffer (25mM TrisHCl 7.5mM; 1M NaCl, 1.8mM MgCl2; 0.5mM DTT and 0.1% NP-40) and one time with buffer B. After the washing steps the beads were magnetically pelleted, the supernatant was discarded and the beads were resuspended in 20mM Tris pH7.5. While on the beads, samples were end-repaired by the T4 PNK enzyme (#M0201L, NEB) following manufacturer instructions. After end-repairing, beads were resuspended in 50ul of proteinase buffer plus 5ul of proteinase K (#3115828001, Roche) and the samples were incubated for 1h at 37ºC with gentle rotation. The RNA from the IPs was purified by adding to the beads 200ul of RNA lysis buffer. Beads were magnetically pelleted and the supernatants were transferred to new tubes. 250ul of ethanol and 25ul of Magbeads (#D4100, Zymo Research) were added per sample. Samples were mixed and incubated for 15 minutes at room temperature with gentle rotation. Beads were magnetically pelleted and washed sequentially with MagBead DNA/RNA Wash1 buffer (#R2130-1, Zymo Research) and MagBead DNA/RNA Wash 2 buffer (#R2130-2, Zymo Research). After magnetically pelleted, beads were resuspended in 30ul of H2O. Samples were incubated at 37ºC for 15 minutes and after magnetically pelleting the beads, the eluted RNA was finally transferred to a new tube.

The saved 100ul of fragmented 2C-RNA for the inputs were processed in parallel during the 2h incubation period of the immunoprecipitation. Samples were processed for a second 2C extraction using a Zymo-Spin IC micro column (#C1004, Zymo Research) and 3ug of the resulting RNA were end-repaired, in a final volume of 50ul, by the T4 PNK enzyme (#M0201L, NEB) following manufacturer instructions. To eliminate the T4 PNK enzyme and buffers, the RNA was later purified by adding 200ul of RNA lysis buffer, after mixing and addition of 250ul of Ethanol, 25 ul of Magbeads were added to each samples. Samples were incubated for 15 minutes at room temperature and gentle rotation. Samples were washed with MagBead DNA/RNA wash buffer 1 and 2 as previously described and RNA was finally eluted in 30ul of H2O. After RNA purification, 1ug of input RNA was subjected to rRNA depletion with the Ribo-Zero Glod Yeast kit (MRZY1324, Illumina) following manufacturer instructions.

RNA purified from the IPs and ribodepleted RNA from the input samples were processed for library preparation with the Nextflex Small RNA kit v3 (#NOVA-5132-06, PerkinElmer), following manufacturer instructions. Libraries from three biological replicates of inputs and IPs were pooled together and sequenced on a NextSeq500 (Illumina) instrument on an 80 pair end run.

### Sequencing informatics analysis

Reads were trimmed with Cutadapt (v2.3) and sequencing quality was inspected with FastQC. Novoalign (v3.07.01) was used to map to the yeast genome (sac3). Gene counts were summarized with featureCounts (v1.6.4). DESeq2 (Love et al.2014) with IHW (Ignatiadis et al.2016) for multiple hypothesis correction was used to determine significantly enriched RNAs in IP samples vs corresponding input controls (adjusted p-value < 0.5; log2 fold-change > 1). Transcriptome coverage plots were generated with bamCompare, computeMatrix and plotProfile functions from DeepTools (Ramírez et al.2014). CSAW (Lun & Smyth2016) was used to detect significantly enriched regions in the IP samples compared to the input controls.

### Fractionation of 2C-RNA in long and short RNA molecules

Four volumes of RNA lysis buffer were added to 100ug of 2C total RNA extracted following the standard 2C method. After mixing, ethanol was added to a final concentation of 33% (vol/vol). Samples we mixed by gentle vortexing and later added to a Zymo-spin IIICG (#C1006, Zymo Research) column. However and as described before, the procedure can be up or downscaled at will. Under these conditions, the column only retains RNA molecules longer than 200nt. The flow through, containing the short RNA fraction, is transferred to a new tube. Ethanol to a final concentration of 65% (vol/vol) is added; samples were mixed and added to a second silica column (mini or micro column), which will now retain the short RNAs. Both sets of columns, containing separately the long and short RNA fractions were loaded with 400ul of RNA Prewash buffer and spun at 10000g for 30 seconds. The flowthrough was discarded and the columns were washed with 700 ul of RNA wash buffer. Columns were centrifuged at 10000g for 30 seconds, the flowthrough discarded and loaded again with 400 ul of RNA wash buffer. Columns were centrifuged at 10000g for 2 minutes to eliminate any residual ethanol. After the centrifugation, the columns were transferred to new collections tubes and RNA was finally eluted in 50ul of H20 by centrifugation at 16000g for 1 minute. To evaluate the performance of the 2C-RNA fractionation, 1 ul of each sample was assessed with Bioanalyzer RNA nanochips (#5067-1511; Agilent) (Schroeder et al.2006). In addition, to accurately analyze the 2C-short RNA fraction, 1ul of the short RNA fraction was also run on a Bioanalyzer Small RNA chip (#5067-1548; Agilent).

### 2C small non-conding RNA interactome capture (snRIC_2C_)

An untagged WT strain was grown in YPD, UV crosslinked and lysed as previously described. Corresponding non-irradiated cells were processed in parallel as negative controls. A first round of 2C total RNA extractions was performed as previously described and 2 vials containing 550 ug of 2C RNA per sample were DNase I treated at 37ºC for 30 minutes in a final volume of 1ml. One set of DNAse I treated RNA was used to purify a second round of 2C total RNA while the second set was used to purify a fraction of 2C short RNA. For the purification of the second round of 2C total RNA, 4ml of RNA lysis buffer were added to the DNase I treated RNA, the samples were gently mixed by vortexing and 5 ml of ethanol, to a final concentration of 50% were added. The samples were mixed again and added to a Zymo-Spin VI maxi column (#C1013, Zymo Research) inserted on a 50ml polypropylene centrifuge tube. Samples were centrifuged at 3000g for 5 minutes and the supernatant was discarded. 4ml of RNA prewash buffer were added and the column was centrifuged again at 3000g for 5 minutes. Later the column was washed twice by centrifugation with 5ml of RNA wash buffer and finally eluted with 2ml of H2O.

For the purification of the second round of 2C short RNA, 2 ml of a 1:1 mixture of RNA lysis buffer and ethanol were added per DNase I treated sample. The samples were mixed and loaded on a Zymo-spin VI maxi column sitting on a 50 ml Falcon tube. Samples were centrifuged at 3000g for 5 minutes, and 3ml of ethanol were added to the flow through, which contained the short RNA fraction. Samples were mixed and loaded on a Zymo-spin IIICG mini column and centrifuged at 10000g for 30 seconds. The flow through was discarded and 400 ul of RNA prewash buffer were added to the column and the samples were centrifuged at 10000g for 30 seconds. Samples were washed two times sequentially by centrifugation with 700 and 400 ul of wash buffer respectively. Finally, 100 ul of H20 were added to the column and the 2C short RNA fraction was eluted by centrifugation at 16000g for 1 minute.

After the second round of 2C, total and short RNA fractions were quantified in Nanodrop 1000 (Thermo Fisher Scientific) and 35ug of each fraction were RNase I (#AM2295; Ambion) digested in 10mM Tris-HCl pH7.5 and 100mM NaCl for 30 minutes at 37ºC. The resulting RNase I treated samples were processed for TMT 6plex labelling as described above for the RIC_2C_ experiment with the exception that 6 fractions per sample were obtained after TMT 6plex labelling. Mass spectrometry data acquisition was done similarly to the RIC_2C_ experiment described above with the exception of the gradient used. In snRIC_2C_, peptides were eluted via the analytical column running solvent A (0.1% formic acid in water, 3% DMSO) with a constant flow of 0.3 µL/min, with increasing percentage of solvent B (0.1% formic acid in acetonitrile, 3% DMSO) from 2% to 8% in 6 min, then 8% to 28% for a further 42 min, in another 5 min. from 28% to 40%, followed by an increase of B from 40-80% for 4 min. and a re-equilibration back to 2% B for 4 min. The outlet of the analytical column was coupled directly to an Orbitrap Fusion™ Lumos™ Tribrid™ Mass Spectrometer (Thermo) using the Nanospray Flex™ ion source in positive ion mode.

### Carbon source depedent tRNA engagement of GAPDH

Precultures of WT and Maf1Δ cells were grown overnight in YPD (2% glucose), YPG (3% glycerol) or YPE (3% ethanol) media. Next day, aliquots were used to start 250ml cultures at an O.D.600= 0.1, and grown until mid-log phase (O.D.600≈ 0.8). Cells were collected and UV-crosslinked, as described before, with 3J/cm2 of UV light at 254nm. After lysis, a 2C total RNA extraction was performed from 1mg of protein lysate and the resulting 2C RNA was quantified in nanodrop. 20ug of 2C RNA from two biological replicates were run on a SDS-PAGE, blotted to a nitrocellulose membrane and probed against GAPDH and ATP5b antibodies.

